## Supplementary figures and images for "Emergent dynamics of adult stem cell lineages from single nucleus and single cell RNA-Seq of *Drosophila* testes"

### Figure 1 - figure supplement 1

#### Figure 1 - figure supplement 1

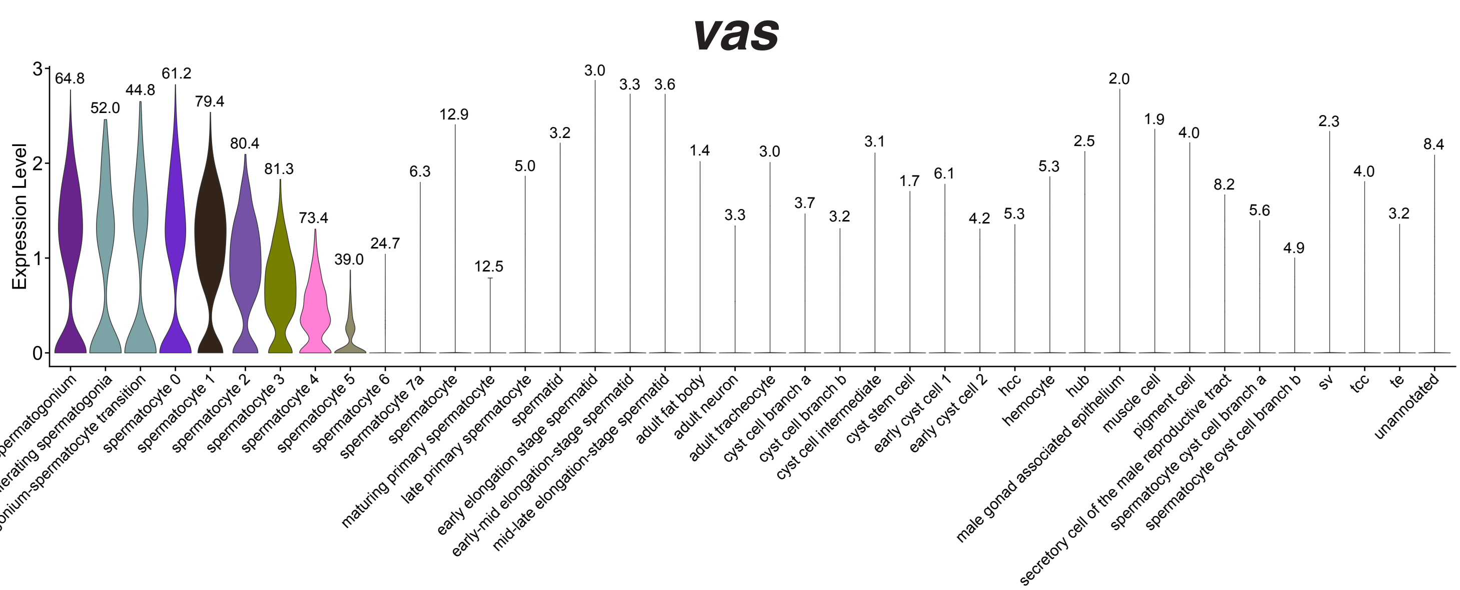

### Figure 2 - figure supplement 1

#### Figure 2 - figure supplement 1

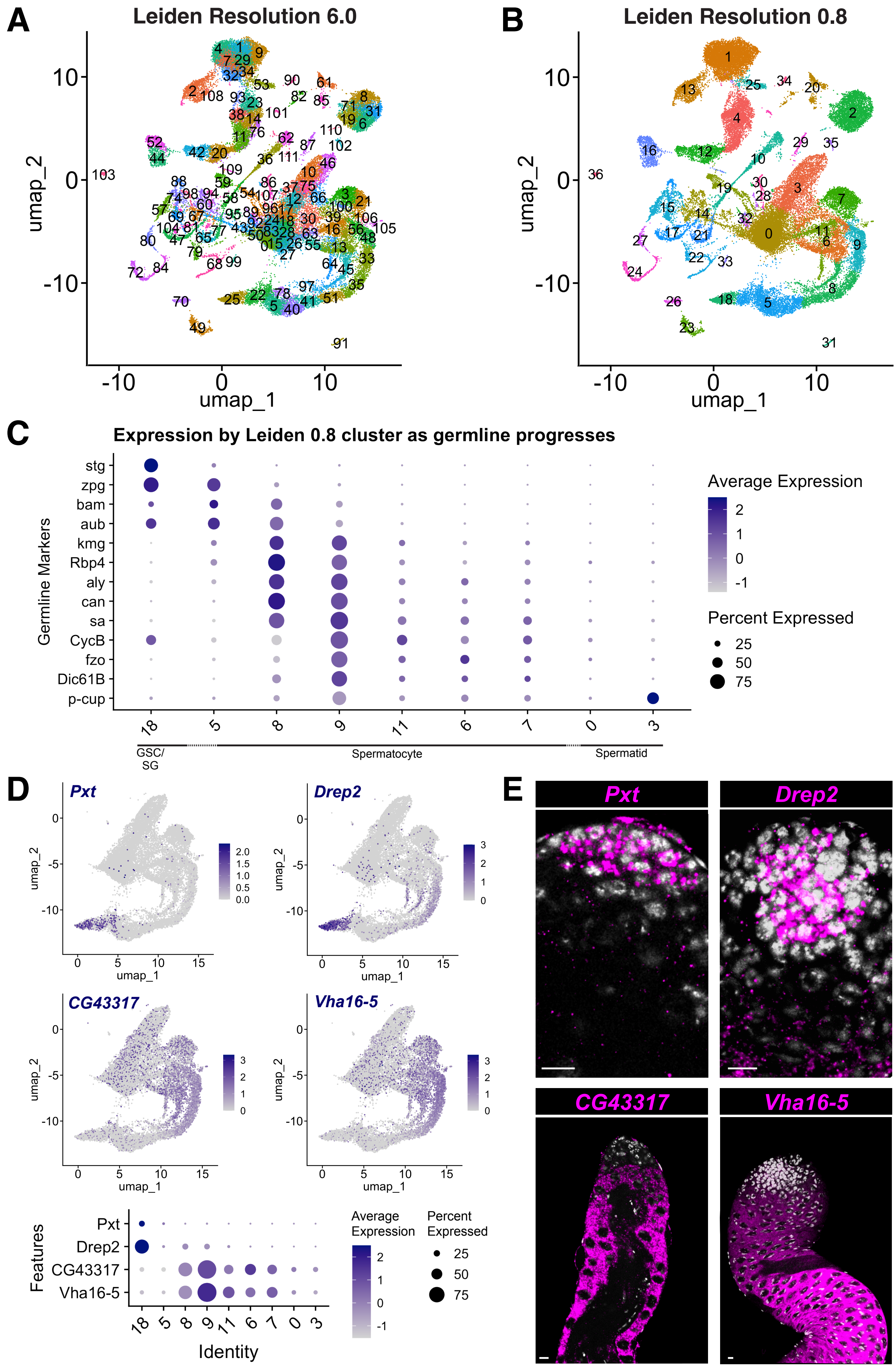

### Figure 3 - figure supplement 1

#### Figure 3 - figure supplement 1

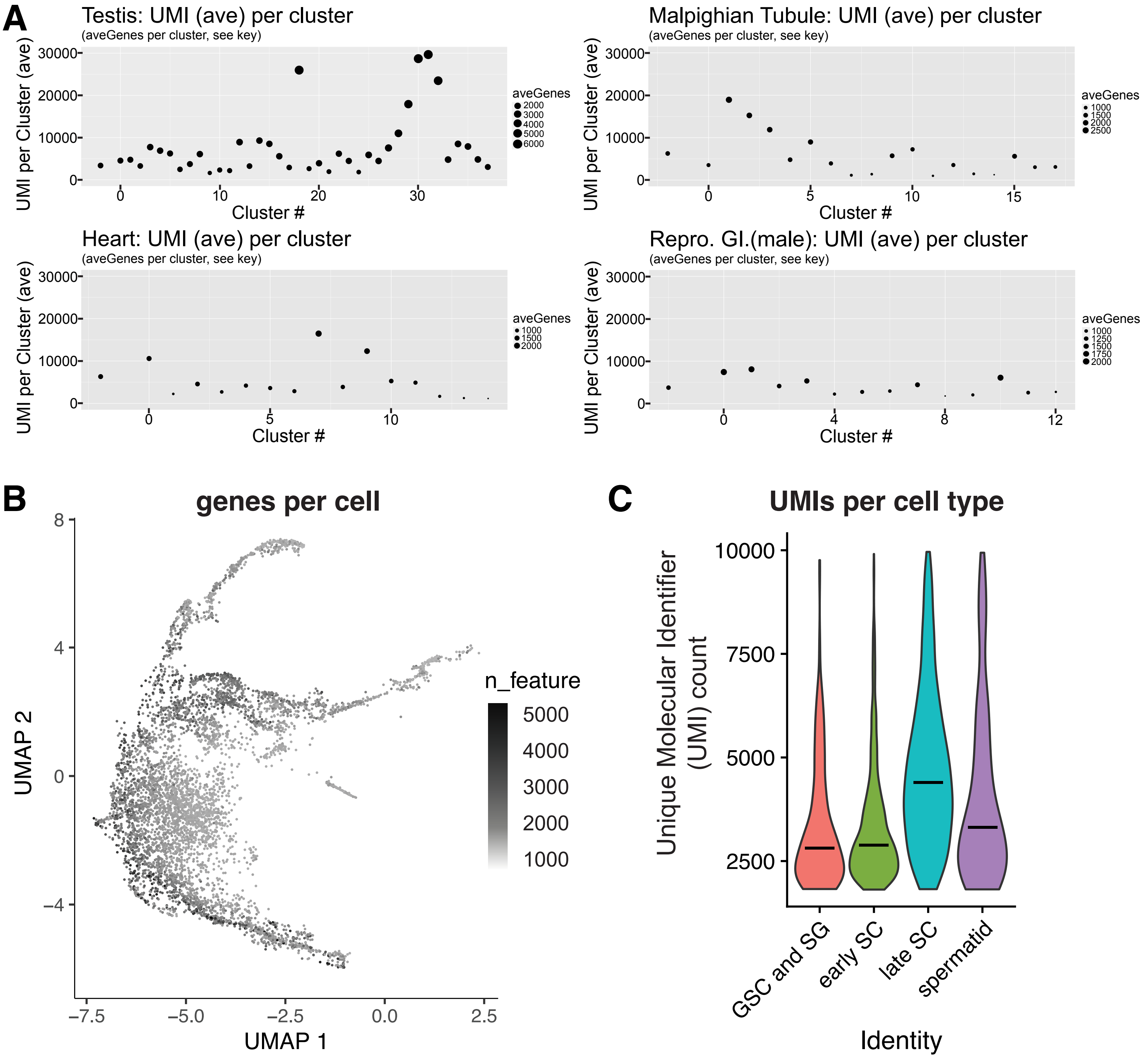

### Figure 3 - figure supplement 2

#### Figure 3 - figure supplement 2

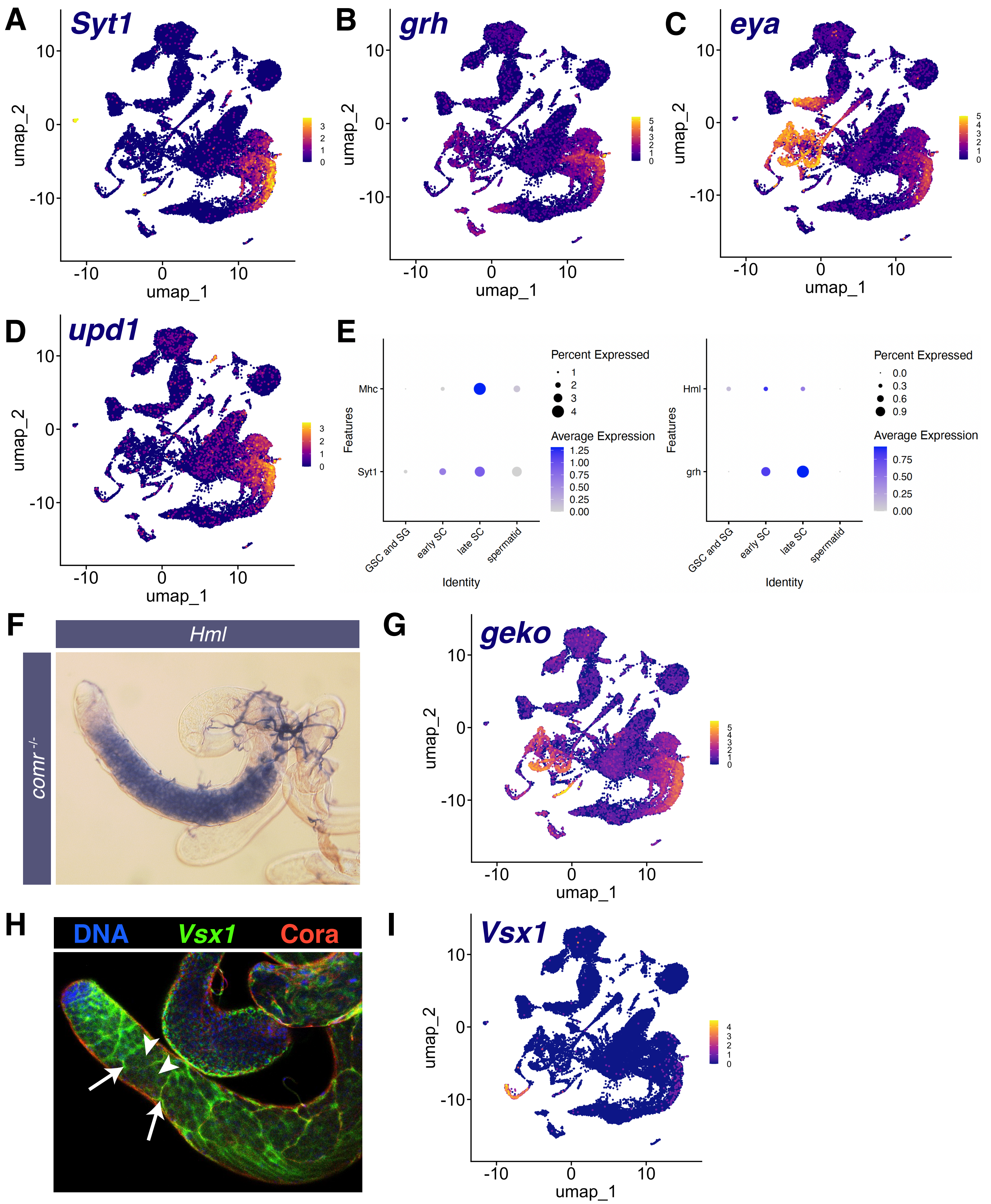

### Figure 4 - figure supplement 1

**Figure 4 - figure supplement 1**

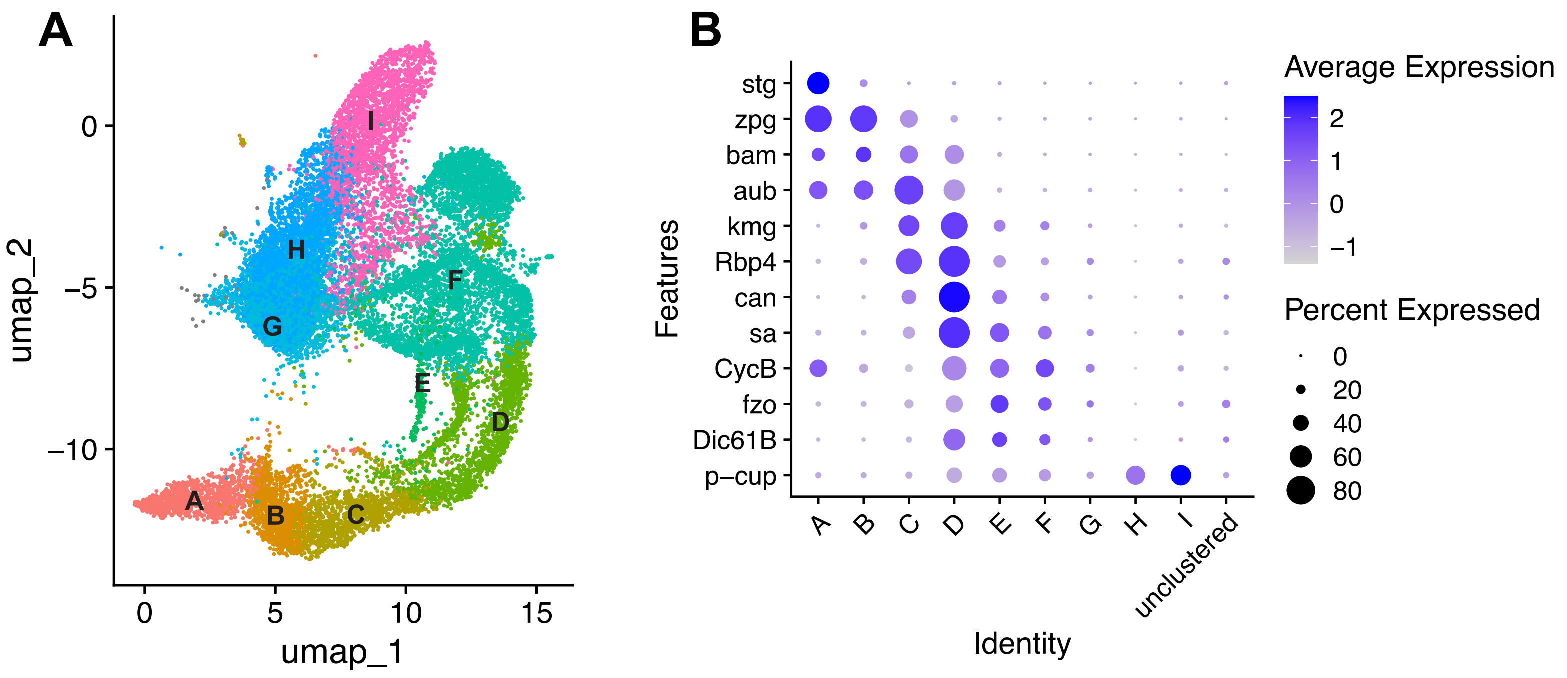

### Figure 5 - figure supplement 1

#### Figure 5 - figure supplement 1

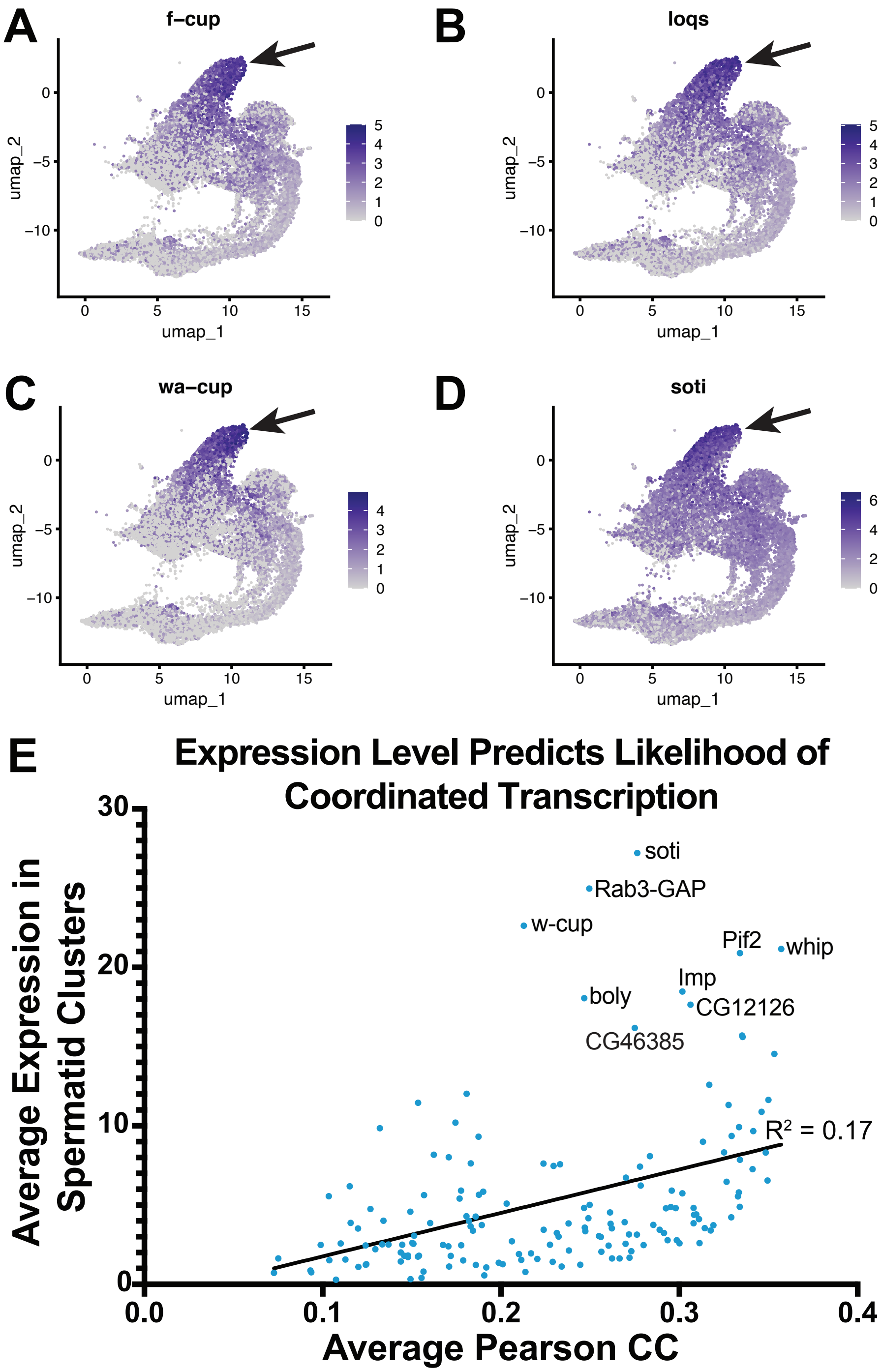

### Figure 6 - figure supplement 1

#### Figure 6 – figure supplement 1

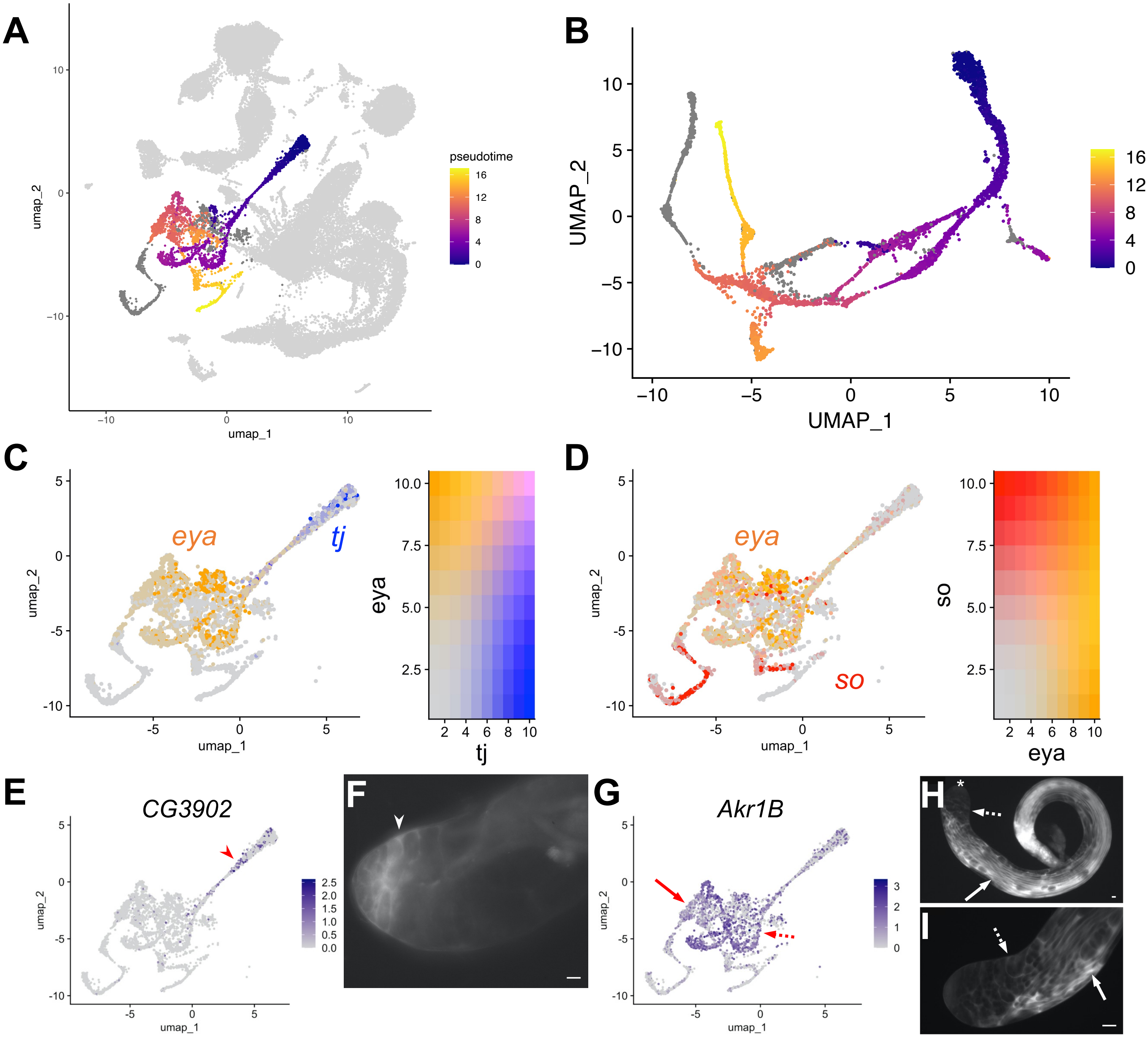

### Figure 7 - figure supplement 1

#### Figure 7 - figure supplement 1

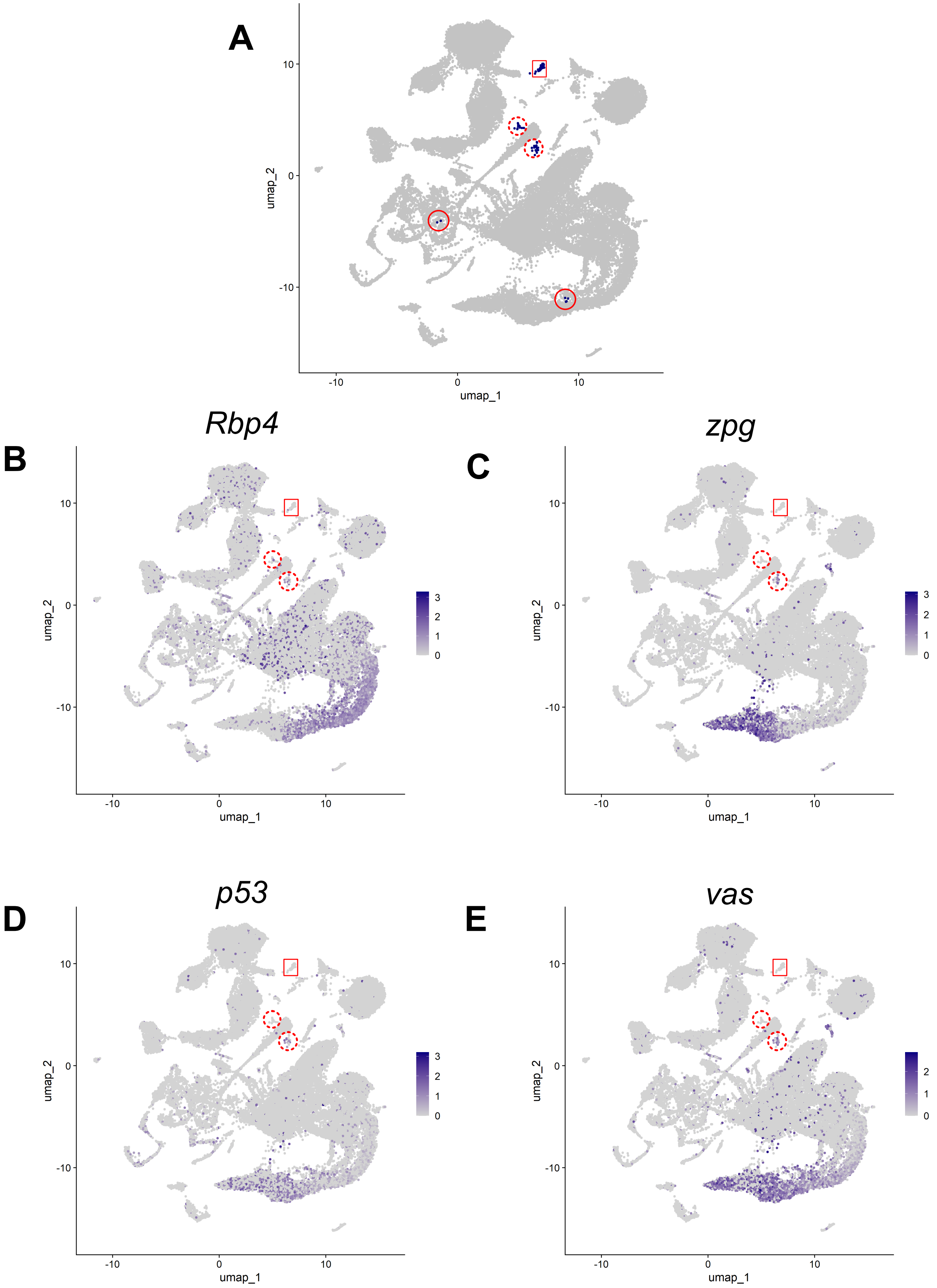

### Figure 7 - figure supplement 2

#### Figure 7 - figure supplement 2

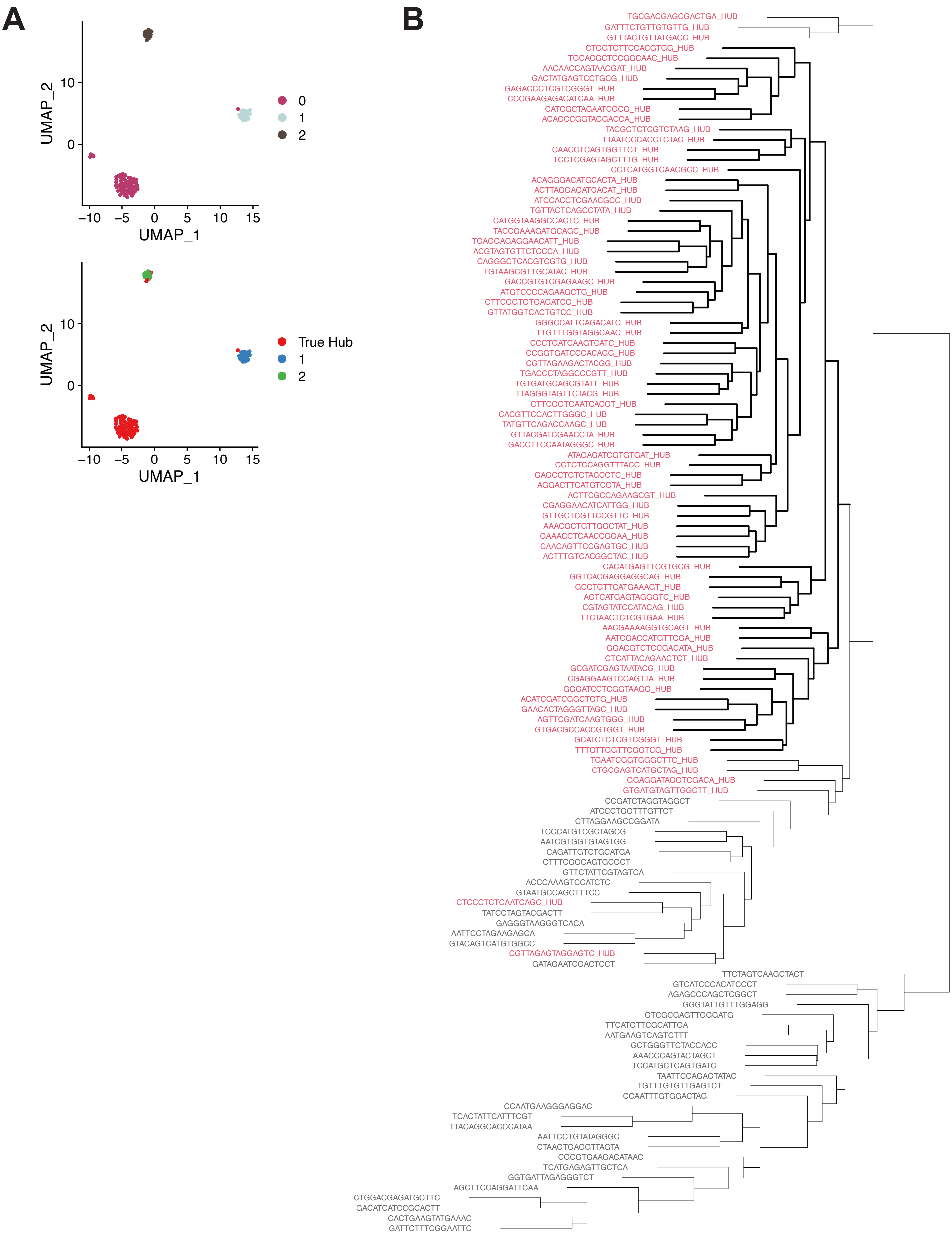

### Figure 8 - figure supplement 1

#### Figure 8 – figure supplement 1

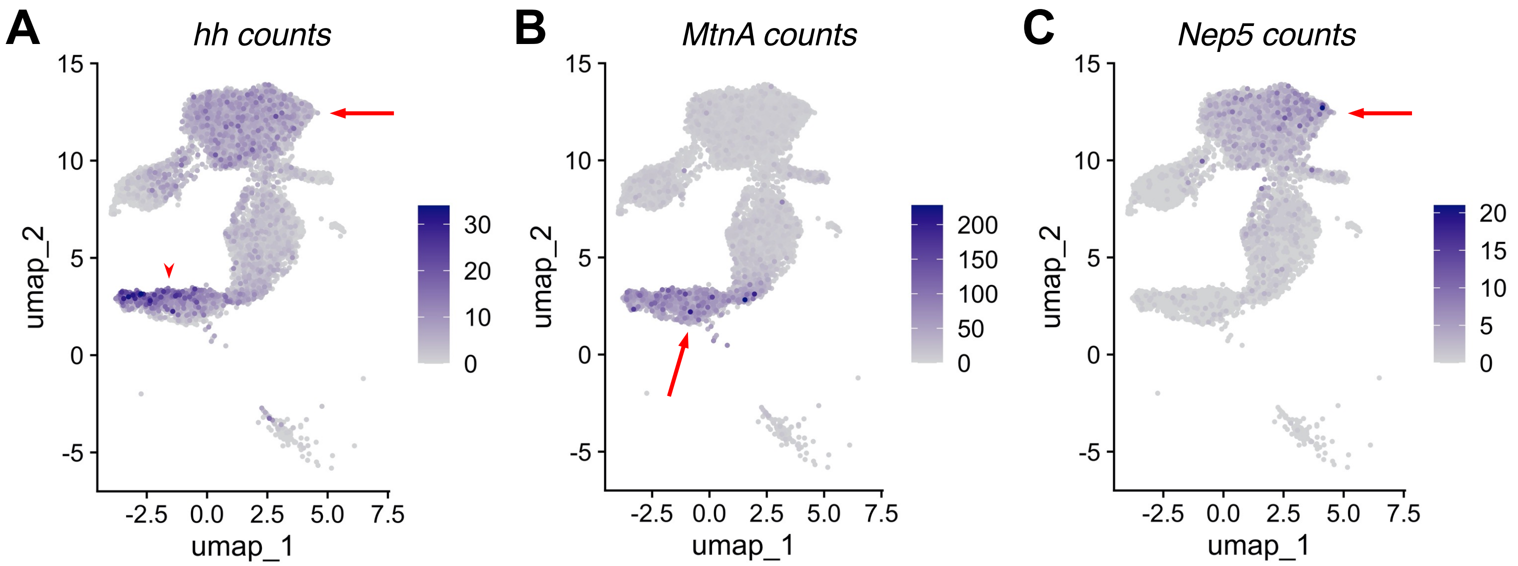
